# Leveraging Machine Learning and AlphaFold2 Steering to Discover State-Specific Inhibitors Across the Kinome

**DOI:** 10.1101/2024.08.16.608358

**Authors:** Francesco Trozzi, Oanh Tran, Carmen Al Masri, Shu-Hang Lin, Balaguru Ravikumar, Rayees Rahman

**Affiliations:** Harmonic Discovery Inc. New York, NY, US; Department of Physics and Astronomy, University of California Irvine, Irvine, CA, US; Department of Chemical Engineering, University of Michigan Ann Arbor, Ann Arbor, MI, US

## Abstract

Protein kinases are structurally dynamic proteins that control downstream signaling cascades by phosphorylating their substrates. Protein kinases regulate their function by adopting several conformational states in their active site determined by the movements of several motifs such as the αC-Helix, DFG residues and the activation loop. Each conformational state represents a distinct physicochemical environment that accepts or precludes ligand binding. However, most of the kinome have not been crystalized across these possible conformational states. It has been shown that shallow Multiple Sequence Alignments (MSA) can enable AlphaFold2 (AF2) to model kinases in alternative conformations. However, it is unclear if these models can be leveraged for structure-based drug discovery. Additionally, there are several machine learning tools to predict protein-ligand interactions based on ligand chemotype and binding pocket properties, but these models cannot be used to identify ligands with clear state specificity. Here, we first present an approach called AlphaFold2 Steering (AF2-Steering), a systematic methodology to direct AF2 to sample kinases in the active and inactive conformations. We use our approach to model the protein kinome in precise conformational states. We demonstrate the utility of these AF2-steered kinase models by employing them in a prospective virtual screening study that integrates machine learning with docking to find state specific inhibitors for well-studied and dark kinases that lack structures in the active conformational state. We then experimentally validate the hits, an essential step often overlooked, and later experimentally confirm the conformation-specificity of the ligands identified for FLT3, a protein kinase that currently lacks an active state crystal structure. Against a strict binding criterion of at least 1μM Kd, our modelled structures achieved an overall hit rate of 53%. We also confirm the conformation-specificity of 4/7 FLT3 ligands, thus demonstrating the value of MSA-steered AF2 modelled kinase structures combined with machine learning and docking to guide conformation-specific kinase drug discovery.

## Introduction

Protein kinases are a diverse group of enzymes that play crucial roles in cell signaling and regulation, making them essential for various biological processes such as cell growth, metabolism, and differentiation^1^. Kinases catalyze the phosphorylation of their downstream substrates, which often serves as a molecular switch, altering the activity, localization, or interaction of the target protein and thereby regulating cellular functions^2^. Due to their ubiquitous presence in biological processes, protein kinases are linked to many different pathological pathways^3–5^, making them a prime target for drug discovery^3,6^.

In modern drug discovery, the availability of protein structures allows scientists to investigate the complementarity between protein targets and small molecules therapeutics at a molecular level. Protein kinases can adopt several pharmacologically relevant conformational states of the active site based on the conformations adopted by the DFG and the *α*C-Helix motifs^7^. The state where the DFG and the *α*C-Helix are both in the “in” position, is the catalytically active conformation of the kinase binding site, called CIDI. The other states are CODI (*α*C-Helix out, DFG in), CIDO (*α*C-Helix in, DFG out), and CODO (*α*C-Helix out, DFG out)^8^. These last three conformational states represent the inactive conformations of the kinase binding site. The diversity of the active site’s physical-chemical environment across these conformational states allows for different modalities of drug binding, which distinguish the different types of orthosteric kinase inhibitors (Type I, Type I-1/2, Type II).

While most kinases have not been crystalized across multiple conformational states^9^, several methods have emerged to model kinases in these states. Earlier methods focused on homology modeling protocols such as DFGmodel^10^, while more recent approaches aim to generate these protein models by leveraging AF2^11^. For instance, Multiple Sequence Alignment (MSA) subsampling coupled with structural templates has been shown to model 437 protein kinases with the catalytic site and activation loop in the active state^12^, or was able to capture alternative conformations of specific kinases^13^. Pure MSA subsampling approaches have been proposed based on stochastic subsampling, varying the max depth in AF2, MSA in-silico mutagenesis, or clustering^14–18^. While these approaches facilitate the exploration of possible alternative conformations, to recapitulate the conformational distribution, or to predict the effect of point mutations, they may not be applicable to steer the generation to a desired conformational state known *a priori*. To this end, a method was recently proposed by Xie et al. to direct the generation towards specific conformations by systematic optimization of the MSA based on user-provided structural features^19^. However, the value of the kinase models produced using these approaches in structure-based drug discovery (SBDD) is yet to be confirmed. Most approaches substitute experimental identification of prospective ligands with *in silico* retrospective analysis such as docking of known ligands^16,20^. Unfortunately, docking has intrinsic limitations in accurately estimating protein-ligand complementarity^21^, which is why experimentally confirmed hit rates from virtual screening have a large degree of variance and can range from 1 to 40%^21^, with 1 to 5% being the typically accepted average hit rate in the industry. Lastly, while numerous machine learning-based tools have emerged to predict protein-ligand affinity^22^, a few tools have been developed to predict the conformational specificity of an inhibitor, many of which are trained solely on ligand features^23,24^. Hanson et al. demonstrated that the same kinase inhibitor can bind to different kinases in distinct conformational states. Therefore, accurately predicting the conformation-specificity of any given protein-ligand complex is crucial for identifying state-specific inhibitors of kinases^25^.

In this work, we first outline an MSA-subsampling approach for modeling of kinases in diverse conformation states called AF2-Steering. We detail the parameter optimization strategy for MSA generation to steer AF2 to generate high quality kinase structures in CIDI conformation. With our strategy we were able to accurately generate high-quality CIDI structures for a set of benchmark kinases and produce CIDI structures for ∼93% of the kinome. Leveraging the CIDI kinase models produced with AF2-Steering, we applied an *in-silico* drug discovery protocol for the prospective identification of kinase inhibitors, which were then evaluated experimentally. By integrating ML binding prediction and docking, our protocol resulted in a hit rate of 53%. We furthermore confirmed state specificity for the inhibitors we identified for the kinase FLT3, which lacks AF2-modelled or crystal structures in the CIDI conformation. We thus provide an experimental validation of AF2 kinase models generated from MSA subsampling by finding conformation-specific kinase inhibitors and determining their conformational-bias for tested kinases.

## Results

### AF2-Steering protocol and identification of optimal hyperparameters

The overall workflow of AF2-Steering is shown in **Fig. 1A**. Given a query protein kinase domain sequence, we first identified homologous kinases across several sequence similarity thresholds (sim. quant.). We then reviewed the structures available in the Protein Data Bank (PDB) ^26^ for each homologous kinase at each sequence similarity threshold. To create conformation biased MSAs, based on the number of available structures we select sequences that have been solved in the desired conformation state with different thresholds of frequency (cidi quant.). Hence, for a query kinase, this strategy results in an array of custom MSAs of various depths. The conformational states of the resulting AF2 models generated using these custom MSAs were assessed using kinformation^8,27^.

**Figure 1.**
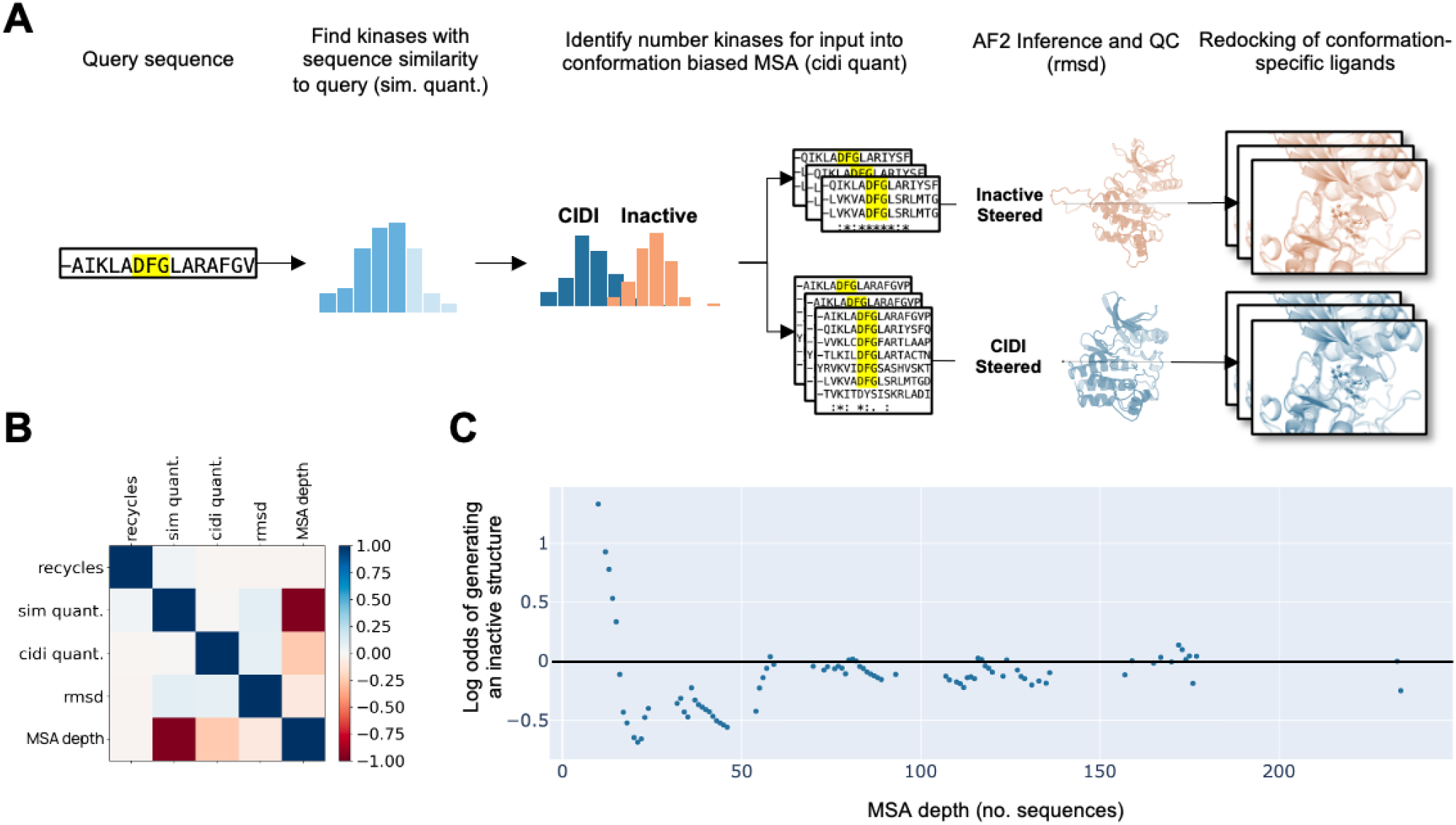
AF2-Steering workflow and hyperparameters evaluation. (A) AF2-Steering protocol. (B) Correlation between hyperparameters and their effect on structural quality. (C) Effect of MSA depth on the probability of producing an inactive conformation.

To evaluate the capability of the AF2-Steering MSA building strategy to effectively steer AF2 generation towards a specific conformation and the structural quality of the generated protein models, we build a benchmark set of kinases. The kinases in the benchmark set have the following properties: i) are not predicted in the target conformation in the AlphaFold DB^28^; ii) have at least a crystallized structure in the targeted conformational state. This benchmark set building strategy allowed us to verify the ability of AF2-Steering to generate structures in the targeted conformational state by comparing them with the AlphaFold DB structures, and to evaluate their structural quality by comparing them to their experimental structure in the same conformational state. The structural quality check was estimated using RMSD as a metric to verify the correctness of overall folding.

By combinatorially modifying the parameters cidi quant., sim. quant., and number of recycling iterations within AF2, we computed a log odds ratio of generating an active or inactive kinase structure and determined the influence of each parameter and their combination in generating the desired conformational state. We observe that certain AF2 parameters like recycling have no effect on the odds of steering the generation towards specific conformational states (**Fig. 1B**). While parameterization of sim. quant. and cidi quant., and in turn MSA depth, have an impact on both the conformation of the generated kinase (**Fig. 1B, C**) and on the quality of the structure generated, as measured by RMSD (**Fig. 1B**). We recapitulate the observation that very shallow MSAs (<20 sequences) are highly likely to generate an inactive kinase conformational state (**Fig. 1B**). We pinpoint, however, that the likelihood of generating an active kinase conformational state peaks at ∼25 sequences, which later diminishes after 50 sequences. Importantly, after 50 sequences, AF2 seems to show less bias towards generating active or inactive conformational states. Furthermore, we observe that deeper MSA depth correlates with lower RMSD (**Fig. 1B**).

### AF2-Steering produces kinases in CIDI conformation

Using this information, we aimed to predict CIDI structures for the entire kinome (**Fig. 2**). As mentioned above, to evaluate the ability of our MSA building strategy to steer predictions towards a target conformation, we benchmarked our models against a set of kinases, found in AlphaFold DB to be predicted in an inactive conformation, but with at least one high-quality structure crystallized in the CIDI conformation for evaluation of structural quality. In this study, we consider a crystal structure high quality if: i) the reported resolution is < 2.5 Å, ii) there are no more than 8 missing residues in the binding site, and iii) is not an atypical kinase. The resulting set is composed of a diverse set of 19 kinases, representative of 6 out 7 kinase groups (Table S1). Our strategy resulted in the production of at least one CIDI structure for all the kinases in the benchmark set (**Fig. 2A**). We further evaluated the percentage of structures predicted in the CIDI conformation using our custom MSA approach vs the default MSA in ColabFold and noticed a 43% increase in generating the CIDI conformation when using our custom steered MSA (Table S1). Our structural analysis of the kinases in the benchmark set to their CIDI reference revealed a high structural similarity to the reference CIDI crystal structures, showcasing the quality of the models (<1.5 Å RMSD, **Fig. 2B**).

**Figure 2.**
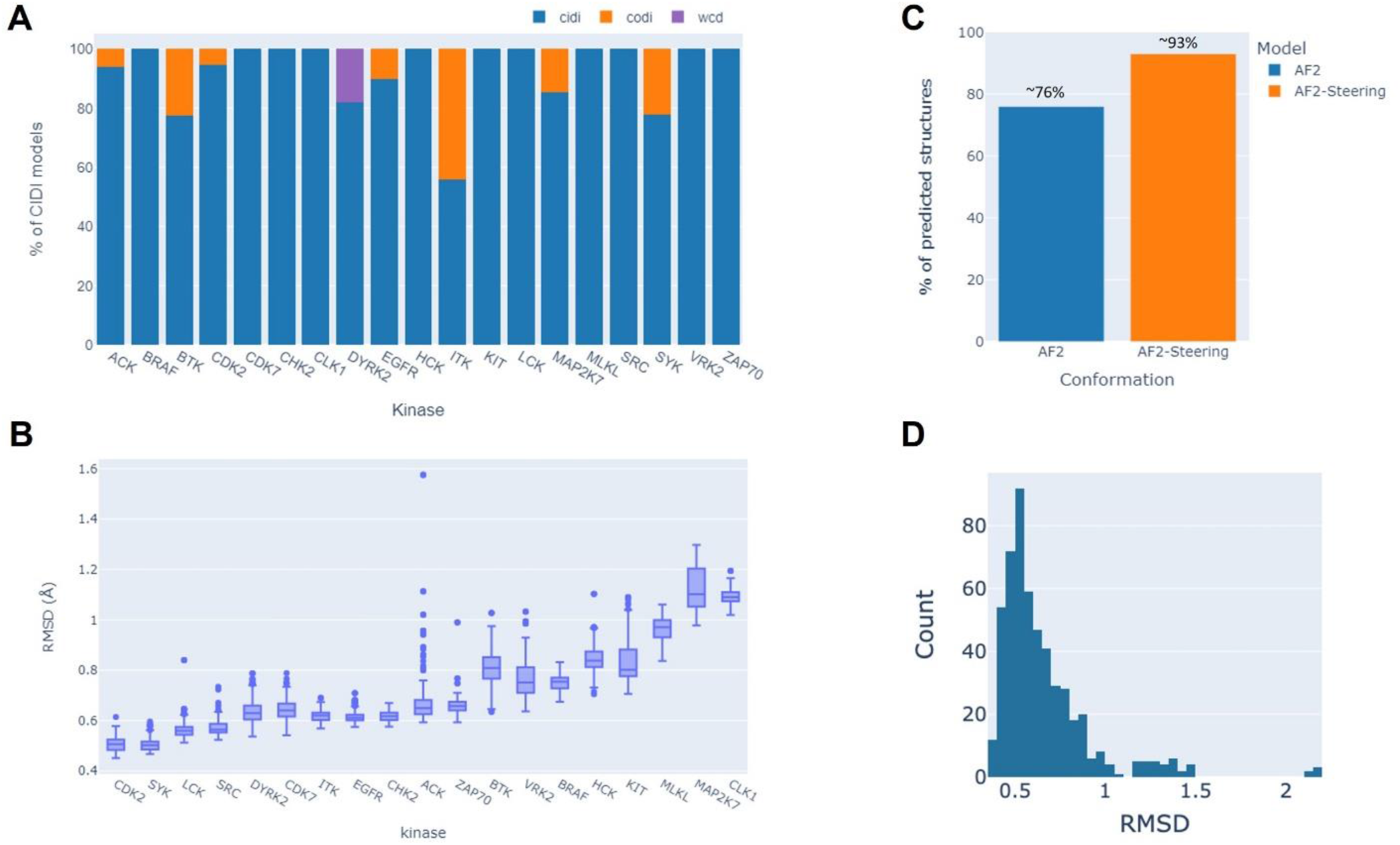
AF2-Steering produces high quality kinase structures in the target conformation. (A) Population of conformations produced by AF2-Steering for a set of benchmark kinases. (B) RMSD of the AF2-Steered kinases in the benchmark set predicted in the target conformation vs their reference crystal structure. (C) Percentage of the kinome produced using AF2-Steering compared AF DB. (D) RMSD distribution of AF2-Steered CIDI structures vs closest CIDI PDB structure.

Once we confirmed that we can steer the conformational sampling of the generated structures towards a target conformation, we moved to the generation of structural models for the entire kinome. Our approach resulted in generating at least one active CIDI conformation for ∼93% of the kinome (460 kinases), compared to ∼76% coverage produced by the AF2 model (**Fig. 2C**). When evaluating these models, we observed that the AF2 Steered models are of high quality, generally having <1 Å RMSD to the nearest kinase crystal structure **Fig. 2D**).

### Experimental validation of AF2-Steered kinase models

While MSA-based methodologies to direct the generation of AF2 towards specific conformational states have been explored in literature, so far, experimental validation demonstrating the usefulness of these models in kinase drug discovery is still warranted. Here, using the structures obtained using our AF2-Steering, we pursued the discovery of inhibitors for 8 protein kinases by implementing a SBDD workflow (**Fig. 3A**). The kinases selected for this study lack active (CIDI) crystal structures. Furthermore, they are either “dark” kinases with little to no known inhibitors, as defined by the dark kinase knowledgebase^29^, or predicted by AF2 to be in their inactive conformation. The dark kinases selected in this study are: BRSK2, CAMK1D, LIMK2, MKNK1. The well studied kinases used for the validation are: DDR2, TYRO3, FLT3, and RIPK1. For each of these kinases we built a conformation-specific library of kinase inhibitors, which lack activity against the target kinases (see Methods section). The libraries were screened using an in-house machine learning-based binding prediction model, described elsewhere^22^. The molecules that had a predicted pIC50 ≥ 7 by the prediction model were then evaluated using docking (see Methods section). While this model can screen for likely binders to the selected kinases, there is no structural information built into the model to estimate the ligand’s conformation-specificity. Post-hoc docking was thus employed to identify ligands that established a type I binding mode within the AF2-Steered models (**Fig. 3**), evaluated versus their matched crystal reference.

**Figure 3.**
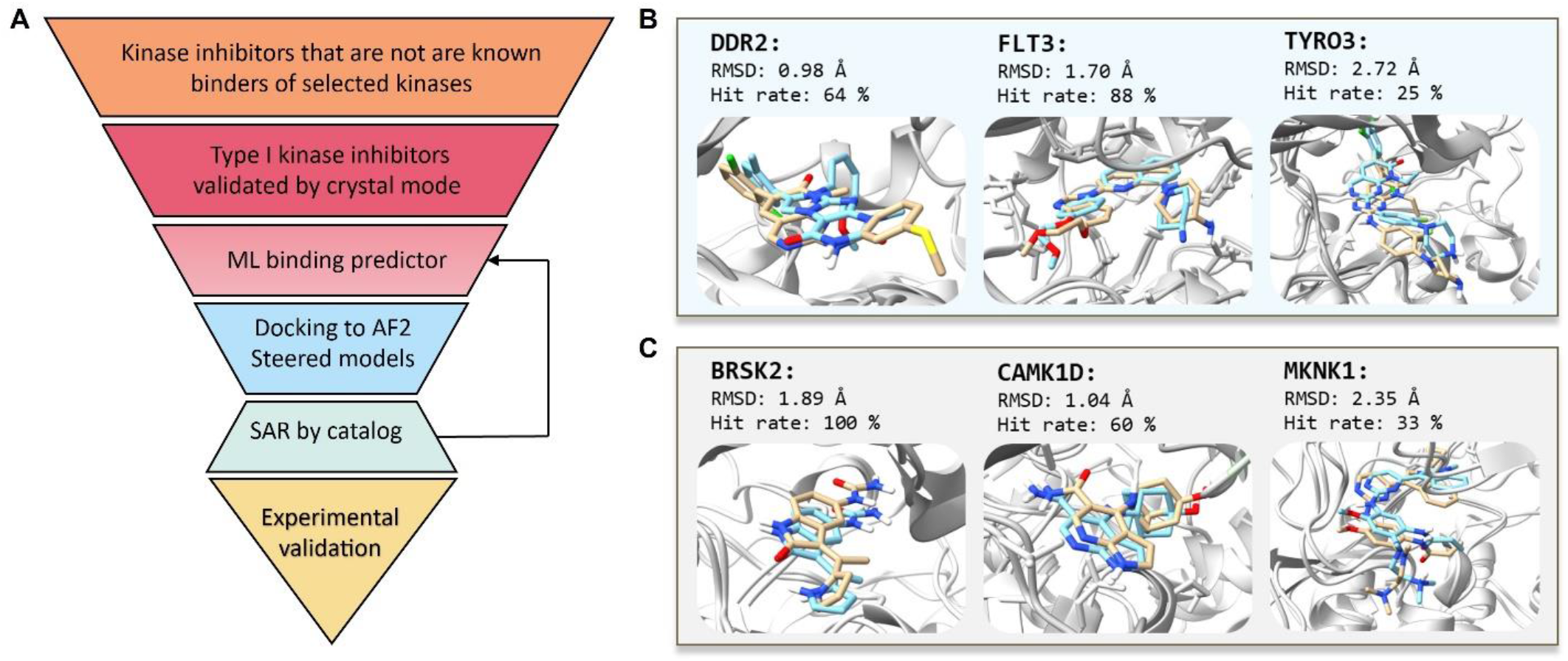
Experimental validation of AF2-Steered structures. A) Structure-based screening workflow for discovering kinase inhibitors. Compilation of docking poses of compounds with high predicted binding affinities to corresponding kinases along with respective hit rate. RMSD comparison to the reference CIDI crystal structure ligand pose for B) well studied kinases and C) dark kinases. Molport IDs for shown hits are: DDR2: Molport-006-168-472, FLT3: Molport-009-680-082, TYRO3: Molport-035-765-968, BRSK2: Molport-046-033-615, CAMK1D: MolPort-044-724-680, MKNK1: Molport-046-862-433.

The resulting hits were evaluated experimentally. The experimental validation using a commercial competitive binding assay (see Method section), was conducted on ligands that exhibited a binding pose RMSD of <3 Å from the matched crystal references (**Fig. 3B, C**). For unavailable ligands, we performed a SAR by catalogue search to identify the closest purchasable ligands, which were then evaluated using the SBDD workflow. The top docking poses of experimentally validated hits for each kinase are shown in **Fig. 3 B, C**. In addition to the low RMSD, the evaluation of these poses shows that these ligands conserve the hinge interactions with the protein, necessary to achieve kinase inhibition, and form the additional interactions in the pocket that are established by their matched reference.

Our SBDD protocol resulted in an overall ∼53% hit rate, with 20 hits out of the 38 kinase-compound pair evaluated. Out of the 8 kinases investigated, we were able to identify potent binders for 6 kinases, out of which 3 are dark kinases (BRSK2, CAMK1D, and MKNK1). The details on the hits, divided per kinase, can be found in Table S2 and S3.

Hanson et al. showed that for the kinase DDR1, a type I ligand may still be able to bind a kinase in the inactive conformation (CIDO, CODI, CODO)^25^. To verify that the hits we identified bind to the CIDI conformation predicted by our AF2-Steering approach, we performed a biochemical binding assay that measured the preferential binding to a specific conformation (see Methods section). For the kinases in our set, this assay was available only for FLT3. This assay is a competitive binding assay which is performed on both the active state of FLT3, which corresponds to the CIDI conformation, and to its autoinhibited state, which corresponds to the inactive conformation of the kinase. Thus, by evaluating the difference in binding of the same ligand to the two states of FLT3, it is possible to evaluate the ligand’s preference for a certain conformational state of the kinase. For FLT3, where CIDI structures were unavailable in both the Protein Data Bank and the AlphaFold DB, out of the 8 hits identified by the SBDD computational protocol, 7 were active with activity at less than 1 μM. We confirmed that 4 out of the 7 compounds preferentially bind to the DFG-in active conformation (**Table 1**). These 4 inhibitors cover 3 distinct chemotypes.

**Table 1:** The binding affinity of hits for FLT3 in the DFG-in and DFG-out (FLT3-autoinhibited) conformation and the respective conformational preferences.

| Compound | FLT3 Kd (nM) | FLT3-autoinhibited Kd (nM) | Preference DFG-in Kd autoinhibited/Kd wt |
| --- | --- | --- | --- |
| 1 | 28 | 10000 | 357 |
| 2 | 0.22 | 17 | 77 |
| 3 | 33 | 2100 | 64 |
| 4 | 110 | 6400 | 58 |
| 5 | 2.6 | 96 | 37 |
| 6 | 460 | 7800 | 17 |
| 7 | 160 | 620 | 3.9 |
| 8 | >10000 | >10000 | 1.0 |

## Discussion

The advent of X-Ray crystallography opened the door to SBDD, which empowered drug discovery with a molecular understanding of protein-ligand interactions. The applicability of SBDD was however limited to the amount of the proteome that was crystallized, or that had close analogues crystallized to perform homology modeling. For example, the evaluation of the number of kinase structures in KLIFS^30^ revealed that 36% of the kinome do not have a solved structure, and only less than 1% have structures across all the key pharmacologically relevant conformations.

Thanks to AF2, SBDD is now, in principle, applicable to the entire proteome- and thus the kinome. However, recent studies have highlighted that for the kinome the conformations produced by AF2 reflect the conformational distribution of the PDB, limiting the conformations that can be targeted for a given kinase^9^. To overcome this issue, multiple methodologies based on AF2 have been developed to increase the variety of conformations produced for a given sequence to produce rarer conformations. While a few of these methodologies can rationally steer the generation towards a specific conformation using MSA subsampling and/or structural templates, to our knowledge an experimental validation to understand the usefulness of these generated structures is still needed.

In this study, we first provided a methodology to steer the generation towards a targeted conformational state called AF2-Steering and, most importantly, we provided much-needed experimental validation of structures generated using this approach. By rigorously evaluating multiple parameters related to MSA generation, we have established a framework for the directed generation of kinase structures in specific conformational states using Colabfold, an open-source implementation of AF2. The capability of AF2-Steering to direct the conformational sampling towards a target conformation was shown through the prediction of CIDI structures for the entire benchmark set. In fact, we observed an overall increase in CIDI prediction to 43% compared to the default ColabFold MSA, and through the prediction of 93% of the kinome in the CIDI conformation, an overall increase of 17% compared to the structures deposited in the AlphaFold DB.

Our experimental validation leveraged the structures generated with AF2-Steering. The validation was performed on a set of challenging kinases for which there was either no structural information in the target conformation, neither experimentally nor in AlphaFold DB, or lacked substantial ligand information. These kinases, called dark kinases, are particularly challenging in screening pipelines that leverage ML binding predictions due to the lack of inhibitor information during the training of these ML binding prediction models. We confirmed the quality of the generated structures by identifying several previously unknown conformation-specific kinase inhibitors for various kinases. Our overall hit rate is 53%, which exceeds the usual hit rate for prospective in-silico virtual screening campaigns^21^. We attribute this high hit rate to several factors: i) the kinase-focused nature of the ligand set, ii) the enrichment of hits with structure-activity relationship (SAR) data from catalogues, and iii) the combination of machine learning (ML) and docking. Notably, the hit rate from this study surpasses that of individual models, where we previously reported that our custom ML binding predictor had a 40% hit rate ^22^ and docking, alone, achieving hit rates in the single digits. We propose that docking complements the high hit rates of our machine learning model by providing information about 3D complementarity, conformation-specificity and necessary binding interactions, thereby effectively reducing the false positive rates associated with purely ML predictions. To this latter point, the result of this study highlights the accuracy of the AF2-steered kinase models in reproducing the pocket physicochemical environment necessary for ligand recognition.

In conclusion, this study confirms the viability of models generated using AF2 MSA-based subsampling and the effectiveness of integrating machine learning approaches with docking for conformation-specific kinase drug discovery.

## Methods

### Custom MSA generation

Kinase-specific MSA were built using the kinome-wide alignment provided by Modi et al.^31^. Using this alignment as input, for each kinase in table S1, multiple MSAs were constructed by varying two parameters, sequence similarity (sim. quant.) and conformational bias (cidi quant.). The sequence similarity between kinases was calculated using the pairwise2 module from the Biopython package^32^. Additionally, the conformational bias of each kinase was assessed by determining the presence of various conformational states in the PDB. Only kinases that have at least one structure in the target conformation were considered for building the MSAs. The inclusion of a sequence in a MSA relative to a target kinase was determined by applying quantile thresholds (0, 0.25, 0.5, 0.75, 0.9) to filter the distribution of these two parameters. For instance, a high threshold of sequence similarity (sim. quant.) and conformational (cidi quant.) would result in only kinases evolutionarily similar to the target kinase and with a majority of PDB structures in the target conformation to be included in the MSA. On the other hand, a low sim. quant. and a high cidi quant., would result in a MSA that includes kinases more distant in the sequence space but that have been crystallized with high frequency in the target conformation. The prediction of the kinase structures was performed using the AF2 implementation in ColabFold^33^. After generation, the conformational state of the kinases in the PDB was evaluated using Kinformation^8^. No relaxation was performed.

### Screening protocol

A proprietary dataset that catalogues known kinase inhibitors was used for the screening. All ligands in the set were featurized using ECFP4 fingerprint and their tanimoto similarity co-efficient (Tc) to ligands crystallized in CIDI kinases were computed using RDKit^34^. Since this dataset is meant to be composed of type-I ligands, ligands that have at least 0.8 Tc to a ligand crystallized in a CIDI structure were retained. The ML binding affinity predictor is an updated version of the top model described in Theisen et al.^22^. Docking was performed using smina^35^, ligands were neutralized, and conformers were built using RDKit^34^. Prior to docking the AF2-steered structures were minimized with a state-specific ligand using MOE^36^ using the Amber10:EHT force field. The RMSD of the maximum common substructure of the docked ligand to the crystal references was calculated using RDKit^34^. Ligands with a docking pose < 3 Å to the reference co-crystalized ligands were selected for experimental validation. Prior to docking and RMSD calculation, all generated AF2-steered structures and crystal structures used as reference were superposed using MOE ^36^. In case a selected ligand was not available for purchase, we searched for available close analogues (> 0.8 Tc). The resulting analogues were filtered according to our screening protocol criteria and, if successful, were purchased and experimentally validated.

### Experimental validation of ligands

The experimental binding affinity was measured with KINOMEscan competitive binding assays (https://www.eurofinsdiscovery.com/solution/kinomescan-technology). Binding to all kinases except for FLT3 was measured using scanELECT. The conformational preference of binding of a molecule for FLT3 was done via the scanMODE assay (https://www.eurofinsdiscovery.com/solution/scanmode). The hit rates for each kinase target were calculated as follows:

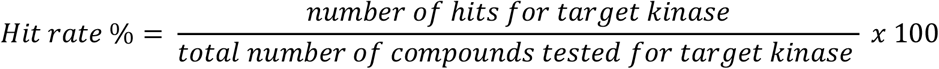

The overall hit rate was calculated as follows:

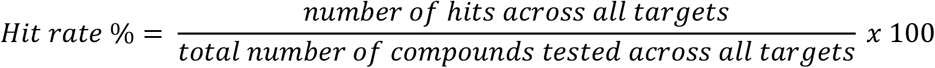

In this study, a hit is defined as > 65 % inhibition @ 1000 nM compound concentration or a Kd < 1000 nM.

### Assays description

#### KINOMEscan

KINOMEscan is a competition binding assay to determine ligand binding to protein kinases. In the KINOMEscan assay, a control ligand is immobilized to a solid support and the protein kinase is tagged with a DNA label. The binding of the test ligands will prevent the kinase from binding to the immobilized ligand. The hits are identified by measuring the amount kinases immobilized, compared to control samples, via qPCR of the DNA label. For more details about the KINOMEscan assay, refer to the original publication^37^.

#### scanELECT

In the present study the concentration of the test compound was 1000nM. The percent of inhibition reported in SI Table 2, is calculated as:

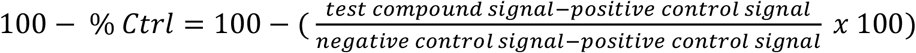

Where:

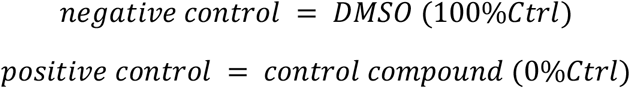

#### scanMODE

The scanMODE is part of the KINOMEscan assay performed on the autoinhibited (inactive) and non-autoinhibited (active) state of FLT3. The binding measurements are done using KdELECT.

The Kd values are determined using the Hill equation to model the dose-response curve.

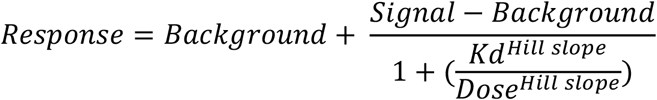

With: *Hill Slope =* −1.

Curves were fitted using a non-linear least square fit with the Levenberg-Marquardt algorithm.

