## Supporting Information for "Leveraging Machine Learning and AlphaFold2 Steering to Discover State-Specific Inhibitors Across the Kinome"

Table S1: Description of kinases in benchmark set and their predicted amount of CIDI conformations.

| Kinase | Kinase group | Kinase Family | Dark | Conformation in Alphafold DB | % of CIDI structures predicted by Colabfold | % of CIDI structures predicted by AF2-Steer | Δ % prediction AF2-Steering - Colabfold default MSA |
|---|---|---|---|---|---|---|---|
| ACK | TK | Ack | No | codi | 40 | 94 | 54 |
| BRAF | TKL | RAF | No | codi | 40 | 100 | 60 |
| BTK | TK | Tec | No | codi | 60 | 77 | 17 |
| CDK2 | CMGC | CDK | No | codi | 0 | 95 | 95 |
| CDK7 | CMGC | CDK | No | codi | 40 | 100 | 60 |
| CHK2 | CAMK | RAD53 | No | codi | 80 | 100 | 20 |
| CLK1 | CMGC | CLK | No | cido | 0 | 100 | 100 |
| DYRK2 | CMGC | DYRK | Yes | wcd | 60 | 82 | 22 |
| EGFR | TK | EGFR | No | codi | 40 | 90 | 50 |
| HCK | TK | Src | No | codi | 100 | 100 | 0 |
| ITK | TK | Tec | No | codi | 0 | 56 | 56 |
| KIT | TK | PDGFR | No | cido | 100 | 100 | 0 |
| LCK | TK | Src | No | codi | 100 | 100 | 0 |
| MAP2K7 | STE | STE7 | No | codi | 0 | 85 | 85 |
| MLKL | TKL | TKL-Unique | No | codi | 0 | 100 | 100 |
| SRC | TK | Src | No | codi | 100 | 100 | 0 |
| SYK | TK | Syk | No | codi | 20 | 78 | 58 |
| VRK2 | CK1 | VRK | Yes | wcd | 100 | 100 | 0 |
| ZAP70 | TK | Syk | No | codi | 60 | 100 | 40 |

Table S2: Details of validated hits across tested kinases.

| Target Kinase | % inhibition @ 1000 nM | Kd (nM) FLT3 | Kd (nM) FLT3 – autoinhibited | % inhibition @ 1000 Nm | PDB ID of Crystal ligand template | RMSD vs template ligand |
|---|---|---|---|---|---|---|
| BRSK2 | 93.2 | na | na | 93.2 | 2pe1 | 1.89 |
| CAMK1D | 70 | Na | na | 70.0 | 6n78 | 2.17 |
| CAMK1D | 98.9 | na | na | 98.9 | 6aaj | 1.04 |
| MKNK1 | 91.5 | na | na | 91.5 | 6lud | 2.35 |
| DDR2 | 100 | na | na | 100 | 5vc5 | 1.77 |
| DDR2 | 100 | na | na | 100 | 5ia3 | 0.99 |
| DDR2 | 98.4 | na | na | 98.4 | 2gqg | 1.77 |
| DDR2 | 99.5 | na | na | 99.5 | 2gqg | 0.97 |
| DDR2 | 100 | na | na | 100 | 2gqg | 1.11 |
| DDR2 | 100 | na | na | 100 | 5vc5 | 1.72 |
| DDR2 | 100 | na | na | 100 | 2gqg | 1.73 |
| DDR2 | 100 | na | na | 100 | 2gqg | 2.67 |
| TYRO3 | 100 | na | na | 100 | 5vee | 0.77 |
| FLT3 | na | 110 | 6400 | na | 4c02 | 0.90 |
| FLT3 | na | 28 | 10000 | na | 4c02 | 2.26 |
| FLT3 | na | 0.22 | 17 | na | 6ble | 1.70 |
| FLT3 | na | 33 | 2100 | na | 6cmj | 1.09 |
| FLT3 | na | 460 | 7800 | na | 1z5m | 2.27 |
| FLT3 | na | 160 | 620 | na | 4c02 | 2.49 |
| FLT3 | Na | 2.6 | 96 | na | 6qav | 1.77 |

| Compound smile |
| --- |
| C\C(=C1\C(=O)Nc2ccc(NC(N)=O)cc12)c1ccc[nH]1 |
| CN(CCC#N)C1CCN(CC1)C(=O)Cn1cc(NC(=O)c2cnn3cccnc23)c(n1)-c1cc(Cl)ccc1OC(F)F |
| NC(=O)c1cnc2[nH]ccc2c1NC1C2CC3CC1CC(O)(C3)C2 |
| COc1cc(N(C)CCN(C)C)c(NC(=O)C=C)cc1Nc1nccc(n1)-c1cn(C2CC2)c2ccccc12 |
| CCN(CC)CCOc1ccc(Nc2ncc3cc(-c4c(Cl)cccc4Cl)c(=O)n(C)c3n2)cc1 |
| CSc1cccc(Nc2ncc3cc(-c4c(Cl)cccc4Cl)c(=O)n(C)c3n2)c1 |
| Cc1nc(Nc2ncc(s2)C(=O)Nc2c(C)cccc2Cl)cc(n1)N1CC[N+]([O-])(CCO)CC1 |
| Cc1nc(Cl)cc(Nc2ncc(s2)C(=O)Nc2c(C)cccc2Cl)n1 |
| Cc1cccc(Cl)c1NC(=O)c1cnc(Nc2cc(nc(C)n2)N2CCNCC2)s1 |
| CCN(CC)CCOc1ccc(Nc2ncc3cc(-c4c(Cl)cccc4Cl)c(=O)n(C)c3n2)cc1 |
| Cc1cccc(Cl)c1NC(=O)c1cnc(Nc2cc(nc(C)n2)N2CCN(CC2)C=O)s1 |
| Cc1nc(Nc2ncc(C(=O)Nc3c(C)cccc3Cl)s2)cc(N2CCN(CCO)CC2)n1 |
| CCn1c2nc(Nc3ccc(N4CCNCC4)c(F)c3)ncc2cc(-c2ccc(Cl)cc2Cl)c1=O |
| COc1ccc(cc1)-c1cnn2cc(cnc12)C#N |
| COc1ccc(cc1)-c1cnn2cc(cnc12)-c1cccc(Cl)c1 |
| COCCOc1ccc2n(cnc2c1)-c1ccc2cccc(N3CCC(N)CC3)c2n1 |
| OC(=O)c1ccc(cc1C1CCCC1)-c1c[nH]c2ncc(cc12)-c1ccccc1 |
| CC(C)(C(N)=O)C(=O)NCCCNc1nc(Nc2cccc(NC(=O)N3CCCC3)c2)ncc1Br |
| c1nn2cc(cnc2c1-c1cccnc1)-c1ccncc1 |
| CN1CCc2cc(Nc3ncc(C4CC4)c(NCCCNC(=O)C4CCC4)n3)ccc2C1 |

Table S3: Hit rate summary.

| Kinase | n. compounds tested | actives | Hit Rate (%) | Dark |
| --- | --- | --- | --- | --- |
| BRSK2 | 1 | 1 | 100 | yes |
| CAMK1D | 5 | 3 | 60 | yes |
| LIMK2 | 3 | 0 | 0 | yes |
| MKNK1 | 3 | 1 | 33 | yes |
| DDR2 | 11 | 7 | 64 | no |
| TYRO3 | 4 | 1 | 25 | no |
| FLT3 | 8 | 7 | 88 | no |
| RIPK1 | 3 | 0 | 0 | no |
